# Characterizing Aerosol Generating Procedures with Background Oriented Schlieren

**DOI:** 10.1101/2022.10.05.511018

**Authors:** N. Scott Howard, Abdulaziz Alrefaie, Nicholas A. Mejia, Tosan Ugbeye, Bryan E. Schmidt

## Abstract

The potential for characterizing aerosol generating procedures (AGPs) using background oriented schlieren (BOS) flow visualization was investigated in two clinical situations. A human-scale BOS system was used on a manikin simulating jet ventilation and extubation. A novel approach to representation of the BOS images using line integral convolution allows direct evaluation of both magnitude and direction of the refractive index gradient field. Plumes issuing from the manikin’s mouth were clearly visualized and characterized in both experiments, and it is recommended that BOS be adapted into a clinical tool for risk evaluation in clinical environments.

## 1 Introduction

The COVID-19 pandemic has brought into sharp focus the risk posed to health care workers by aerosol-generating procedures (AGPs) [1]. AGPs are medical procedures that produce aerosols that can be laden with pathogens if the patient has a viral infection, such as SARS-CoV-2 or influenza. These aerosols pose a significant risk to health care workers, as they can linger in the air for extended periods of time and are small enough to bypass ordinary surgical masks.

The COVID-19 pandemic has exposed significant gaps in our knowledge concerning AGPs [2], specifically: (i) What exactly constitutes an AGP? (ii) How do we characterize AGPs? (iii) How are the risks of a particular AGP quantified? (iv) What strategies are most effective to minimize the risk of pathogen transmission during an AGP?

It is extremely important that these gaps be addressed, not only to protect health care workers during the current pandemic, but also to fight against any severe infectious diseases that may arise in the future. One key question to be addressed in this study is how to best visualize the aerosols generated during AGPs, so that risk can be assessed and mitigation strategies developed.

The risk of AGPs cannot be readily evaluated if the flows containing the aerosols are not understood. Recent studies have shown that airborne viral transmission range is not merely determined by a dichotomy of particle size - “large” or “small”, also known as droplets or aerosols, but rather that there is a continuous dispersion of sizes carried in a multiphase turbulent plume [3,4]. Thus, any activities that increase the velocity or volume of air forced over respiratory mucosa provides increased potential risk which we feel cannot be solely measured with particle counters.

There are several flow visualization techniques developed within the fields of engineering and fluid mechanics, but most are not suitable for use in a clinical setting, and have therefore not been utilized in this context other than for non-human studies. For example, techniques involving light scattering from a laser sheet, such as Mie [5] or Rayleigh scattering [6] would be dangerous to patients and health care workers due to the high power Class IV lasers required. Many techniques are overly intrusive or impractical because of the need to add additional liquid droplets or solid particulate to the flow to enhance light scattering.

Schlieren, on the other hand, is a non-intrusive method for visualizing optical index variations in transparent media [7], and has been successfully employed to visualize the flow produced by coughs, sneezes, and ordinary respiration in laboratory settings to assess the effectiveness of masks in reducing the spread of airborne diseases [8–10]. It is a common tool for the visualization of gaseous flows in the fields of aerospace engineering and fluid mechanics. Schlieren does not require potentially hazardous equipment such as lasers, and it is able to directly visualize refractive index gradients caused by changes in gas composition, aerosols, and/or changes in the gas density via the Gladstone-Dale relation. Conventional schlieren uses a beam of collimated light created by a pair of large focusing lenses or mirrors which is passed through a region of interest with variable optical index. The interested reader is referred to the reference text by Settles [7] for more information.

Conventional schlieren imaging, however, requires a precise optical setup that makes it difficult to employ outside of a laboratory that is specifically designed for schlieren visualization. Furthermore, conventional schlieren is difficult to perform by individuals who are not experienced experts. Additionally, the field of view of a traditional schlieren setup is limited by the diameter of the focusing mirrors or lenses used. Tang et al., for instance, required a focusing mirror with a 1 m diameter to fit the upper torso and head of a human adult in their field of view [8, 9]. Large focusing optics such as these quickly become prohibitively expensive, in addition to being extremely heavy and therefore non-portable.

Background Oriented schlieren (BOS) is a recently developed experimental technique in fluid mechanics for visualizing optical index gradients in variable-density flows that is an alternative to conventional schlieren [11]. The technique works by imaging a patterned background first with no disturbances in the foreground, and then again with a subject in the foreground that causes a distortion to the background pattern, such as the flow of an aerosol. Flows are visualized by computing the apparent displacement of the background pattern between the initial image and subsequent video using a custom image processing algorithm.

While BOS is able to visualize optical index variations non-intrusively like traditional schlieren, it does not suffer from some of its drawbacks [11]. It is easy to set up, easy to use, and the system does not require any large pieces of equipment other than an appropriate background and camera. BOS can therefore provide the ability to visualize the flows of aerosols created by AGPs in a clinical setting and hence enable the unambiguous evaluation of the risks associated with the spread of airborne pathogens. BOS was recently applied by Mok et al. to visualize the plumes exhaled by women during labor and delivery [12]. The overall goal of the present study iss to investigate a flow visualization-based method in a human scaled clinical setting to evaluate risks and develop risk-mitigation strategies for AGPs. Our hypothesis is that background-oriented schlieren (BOS) will be an effective technique for visualizing aerosols produced during AGPs.

## 2 Methods

The experimental setup used for BOS imaging is shown in Fig. 1. The subject of interest, in this case a manikin, is positioned between the camera and patterned background. A camera is focused onto the background, and a reference image *I_r_* is acquired with no disturbances between the camera and the background. A new set of reference images are recorded before the start of each experimental condition. After the reference image is acquired, light rays propagating from the background to the camera are refracted by an angle ε as they pass through a field of varying optical index *n* at the subject caused by an exhaled flow, which is placed a distance *Z_D_* from the background, between the background and the camera. A set of data images *I_d_* (i.e., a video) is then acquired, where the background pattern has been distorted relative to the reference image in each data image. An example raw data image and the corresponding processed BOS image are shown in Fig. 2.

**Fig. 1.**
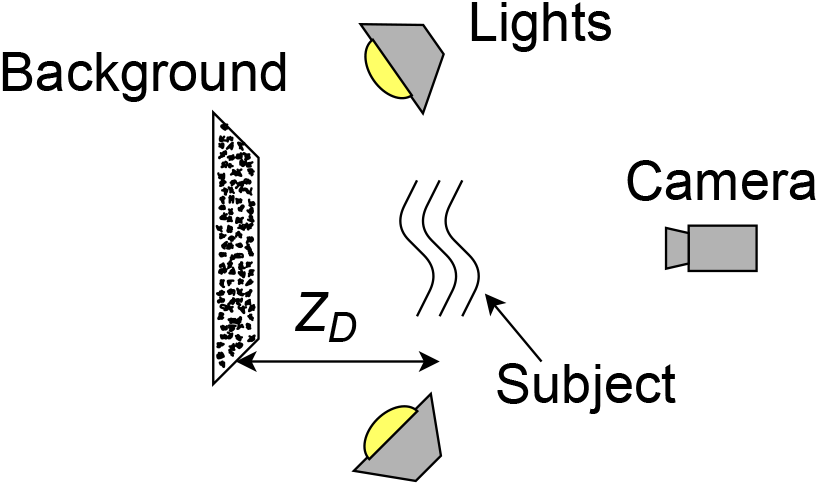
A typical BOS setup, viewed from above. *Z_D_* is the distance from the subject to the background.

**Fig. 2.**
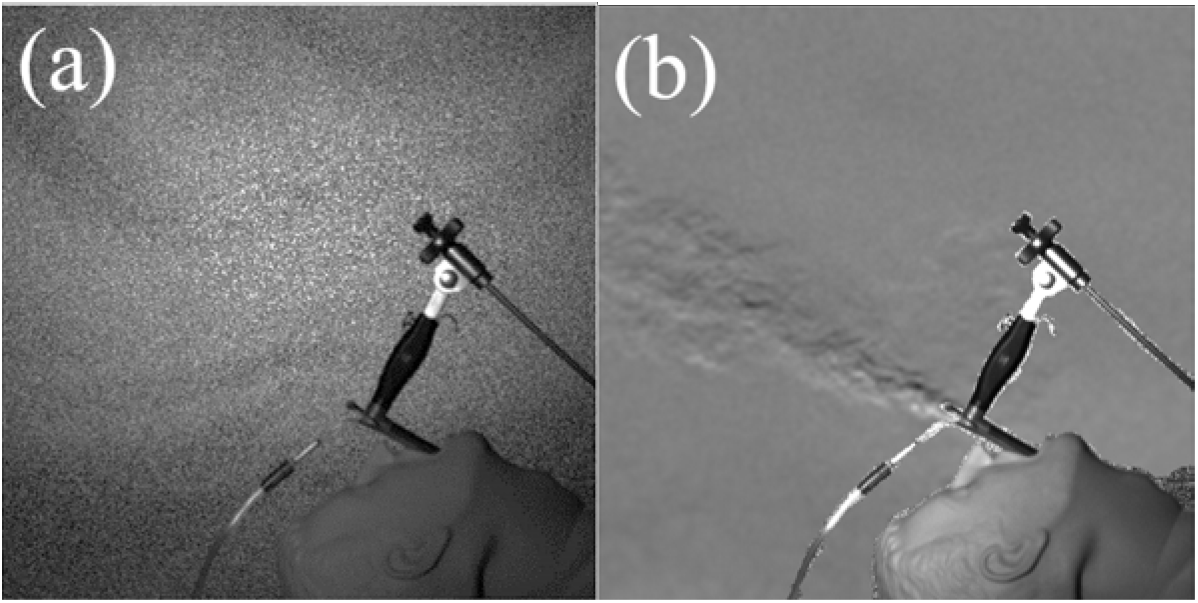
Example BOS images of jet ventilation. (a) Raw BOS image, showing the background pattern. (b) Processed BOS image, showing the aerosol plume issuing from the manikin’s mouth during jet ventilation.

The apparent displacement of each point on the background pattern in the imaging plane of the camera, Δx, is given by

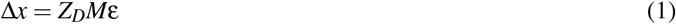

where *M* is the magnification of the camera. The larger Δx is for a given refraction angle ε, the more sensitive the BOS system is. In order to produce the final BOS image, like the one shown in Figure 2b, the data and reference images must be compared using an image processing algorithm to determine the pixel-wise displacement between the reference image and subsequent images. The choice of the algorithm used is crucial, as it has a significant effect on the final resolution and quality of the processed BOS images. In this work, we use the wavelet-based optical flow analysis (wOFA) method developed in [13]. BOS has most often been applied for the visualization of relatively small, laboratory scale test objects, with a maximum dimension on the order of 30–50 cm. For healthcare settings, however, a much larger field of view is required in order to capture human-scale flows, which presents several challenges.

To illustrate these challenges, we can rewrite *M* in Equation (1) using the camera sensor size *h_i_* and the size of the background *h_o_*:

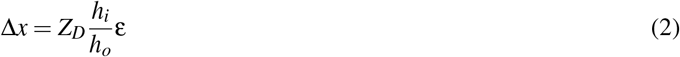

Hence, the sensitivity Δ*x* for a given test subject decreases as the background size *h_o_* increases. Note also that as the size of the subject increases, the size of the background must increase as well. For example, in the experiment shown in Figure 2, the field of view in the plane of the subject was 60 × 60 cm and the corresponding field of view in the background plane was 169 × 169 cm. The full size of the background screen used is 244 × 183 cm. One way to counteract the loss in sensitivity is to increase the distance from subject to background, *Z_D_*, but this in turn causes several additional challenges within the clinical scenarios envisioned, and so is not necessarily a viable solution [11].

These limitations are largely addressed in the present work by the high sensitivity and spatial resolution of the optical flow BOS algorithm. Schmidt and Woike demonstrated that the wOFA method is significantly more sensitive to small displacements than conventional BOS algorithms, by up to a factor of three, and can accurately resolve sub-pixel displacements [13]. This ability to resolve small displacements allows the wOFA algorithm to be applied to BOS images with larger fields of view than would be possible with a conventional algorithm.

BOS has one other advantage over conventional schlieren that is very seldom taken advantage of. Conventional schlieren setups are only capable of visualizing one component of the refractive index field, the one perpendicular to the knife edge. BOS, on the other hand, resolves both components of the displacement vector field between the reference and data images. This additional information is not typically presented in the literature, however, and often only a single component of the displacement field is shown in order to make a more direct comparison to conventional schlieren, as in Fig. 2b.

This may be due in part to the difficulty of representing two-dimensional data, i.e. a vector field, in a simple way that can be interpreted intuitively by the human eye. Laidlaw et al. [14] conducted a quantitative, comparative study to evaluate the effectiveness of various vector field visualization strategies utilizing both expert and non-expert test subjects. They found that line integral convolution (LIC) produced superior results for half of the study criteria, compared to five other visualization strategies. LIC was developed by Cabral and Leedom [15], and has been adapted and used extensively in the fields of fluid mechanics and image processing. Briefly, LIC filters an image containing a randomized texture, often Gaussian white noise, using a two-dimensional vector field that is integrated along its streamlines. This creates an image of textured curves which follow the streamlines of the flow. The resulting image can then be colored to represent the vector magnitude. To the best of the authors’ knowledge, LIC has not been used previously for BOS images, although interestingly, LIC was used to visualize synthetic displacement fields used as known inputs to BOS algorithms by Atcheson et al. [16], but it was not used to visualize the resulting BOS images.

Figure 3 demonstrates the use of LIC on a BOS image from a jet ventilation experiment. Instead of using Gaussian white noise as the input texture for the LIC image, an image of wavelet noise [17] is used. Wavelet noise is multi-scaled, and has a more “natural” appearance than Gaussian white noise, and was therefore judged to be more appealing for visualization purposes. Unlike for the application of LIC to traditional vector fields, the components of the BOS displacement vector field are swapped prior to computing the LIC image. This leads to visualization of isocontours of the refractive index gradient, instead of viewing lines along the gradients. Visualizing the isocontours results in a more compelling and intuitive image, as observed in Fig. 3b. The displacement vector magnitude is represented by coloring the LIC image with a perceptually uniform color map [18]. Compared to visualization of the vertical component of the displacement only, in a way which mimics conventional schlieren images (Fig. 3a), the LIC image is simultaneously more visually appealing and more informative, as it conveys information not only about the concentration of the aerosol in the plume by the magnitude of the displacement, but also reveals turbulent structures along the isocontours of the refractive index gradient. The large-scale, small-magnitude displacements in the regions outside of the plume are due either to motion of the background screen, ambient air currents in the room, or a combination of both.

**Fig. 3.**
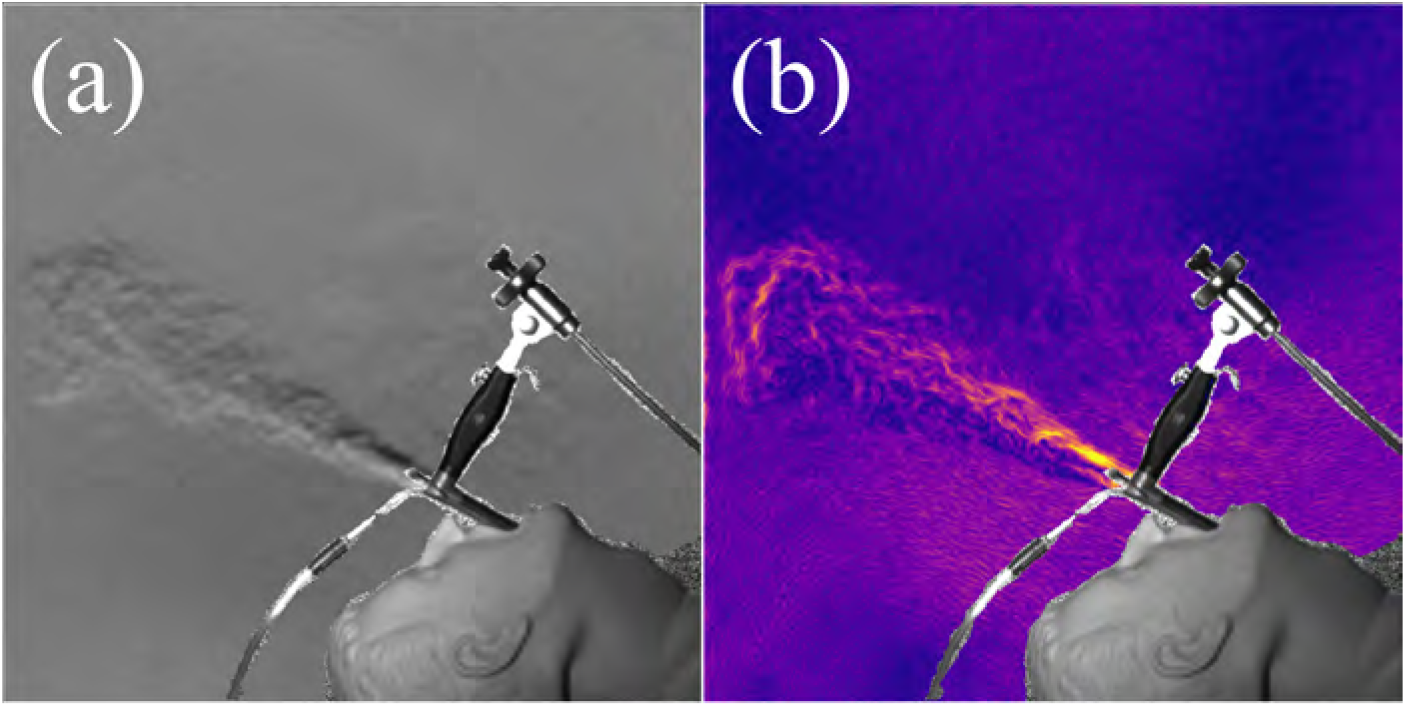
Demonstration of line integral convolution (LIC) for a BOS image. (a) A “traditionally” visualized BOS image, showing the vertical component of the displacement only to imitate a conventional schlieren image. (b) BOS image visualized with LIC, coloration corresponds to the displacement magnitude.

### 2.1 Adapting BOS to a Clinical Setting: Jet Ventilation and Extubation

Jet ventilation was implemented using a Dedo laryngoscope and a manikin with simulated lungs (Laerdal Airway Management Trainer). Extubation was modeled with a similar manikin, and air was supplied to the manikin’s lungs with a manual resuscitator directly connected to the airway. During the extubation trials, a small amount of carbon dioxide (CO_2_), approximately 50% by volume, was introduced in order to increase the variations in the index of refraction and hence boost the signal to noise ratio of the BOS images over air alone. This will likely not be necessary with actual human test subjects, as the temperature and natural CO_2_ and aerosol content of a human breath will provide sufficient image quality [12]. A high-resolution, high speed camera (Photron Fastcam Mini AX100, San Diego, CA) was utilized for the jet ventilation study, and a similar camera (Photron Fastcam Nova S12) was used for the extubation study. In both cases, Photron PFV4 software was used to acquire the images. The high resolution of the cameras (> 1MP) is important, as it increases the sensitivity of a BOS system. From an image processing standpoint, the displacement on the camera sensor is ultimately measured in the number of pixels instead of physical length. Therefore, a sensor with the same size but twice as many pixels in a given direction will be twice as sensitive for BOS.

A Gaussian noise pattern was used as the BOS background, which was printed on a 244 × 183 cm sheet of canvas. The choice of background pattern affects the quality of the processed BOS images, and Gaussian noise has been demonstrated to produce superior results for the wOFA algorithm used in the present study [13]. The canvas sheet was stretched on a collapsible rigid metal frame and secured to the wall behind the subject in order to minimize movement of the background by air currents in the room. This is necessary, because physical motion of the background pattern between the reference and data images will be interpreted by the processing algorithm as optical distortions due to a flow, and hence will be visible. The background screen was illuminated by two flicker-free 75 W white light LED lamps (Nila Varsa Daylight, Altadena, CA).

## 3 Results

### 3.1 Jet Ventilation

The BOS system as described was able to visualize the flows produced by jet ventilation, a specific example of an AGP (see Figs. 2 and 4). Jet ventilation is a technique of apneic ventilation where high pressure flows of oxygen or mixed air are delivered by a variety of techniques into the lungs. Inspiration during jet ventilation is often achieved by insufflation of pressurized gas through a cannula placed within a laryngoscope that enters into the patients mouth and sits directly above the vocal folds. Alternatively, a jet ventilation catheter is inserted below the vocal folds in a fashion similar to tracheal intubation. Expiration is passive as a result of the elastic recoil of the lungs and the chest wall. In our experiments, the plume issuing from the mouth and through the laryngoscope is clearly visible, indicating that BOS is sufficiently sensitive to evaluate the risk of AGPs by characterizing the size and location of the aerosol-laden flow produced by this AGP. These initial experiments were helpful to define an optimal arrangement of the subject, background screen, and camera to maximize the SNR within the constraints of the room dimensions. Since background patterns can be easily changed using a computer, we subsequently optimized the background pattern to the 244 × 183 cm screen. This portable, large field of view BOS system will allow for new studies of a plethora of flows of engineering and medical importance.

Before processing the reference and data images using the wOFA BOS software [13], portions of the image where the background is not visible were masked using a program with a graphical user interface implemented in MATLAB. These regions include the manikin head, the laryngoscope, and the ventilation tube. Not using a mask can cause erroneous displacements to be calculated by the algorithm in these parts of the images, which can influence the computations at nearby pixels due to the regularization used in the wOFA code. After the BOS images were processed, the regions of the original images that were masked out are superimposed on the schlieren fields, to produce the images shown in Fig. 4. Figure 4 shows the vertical component of the 2D displacement field, and maximum displacements of 1.5–2 pixels were observed in these experiments.

**Fig. 4.**
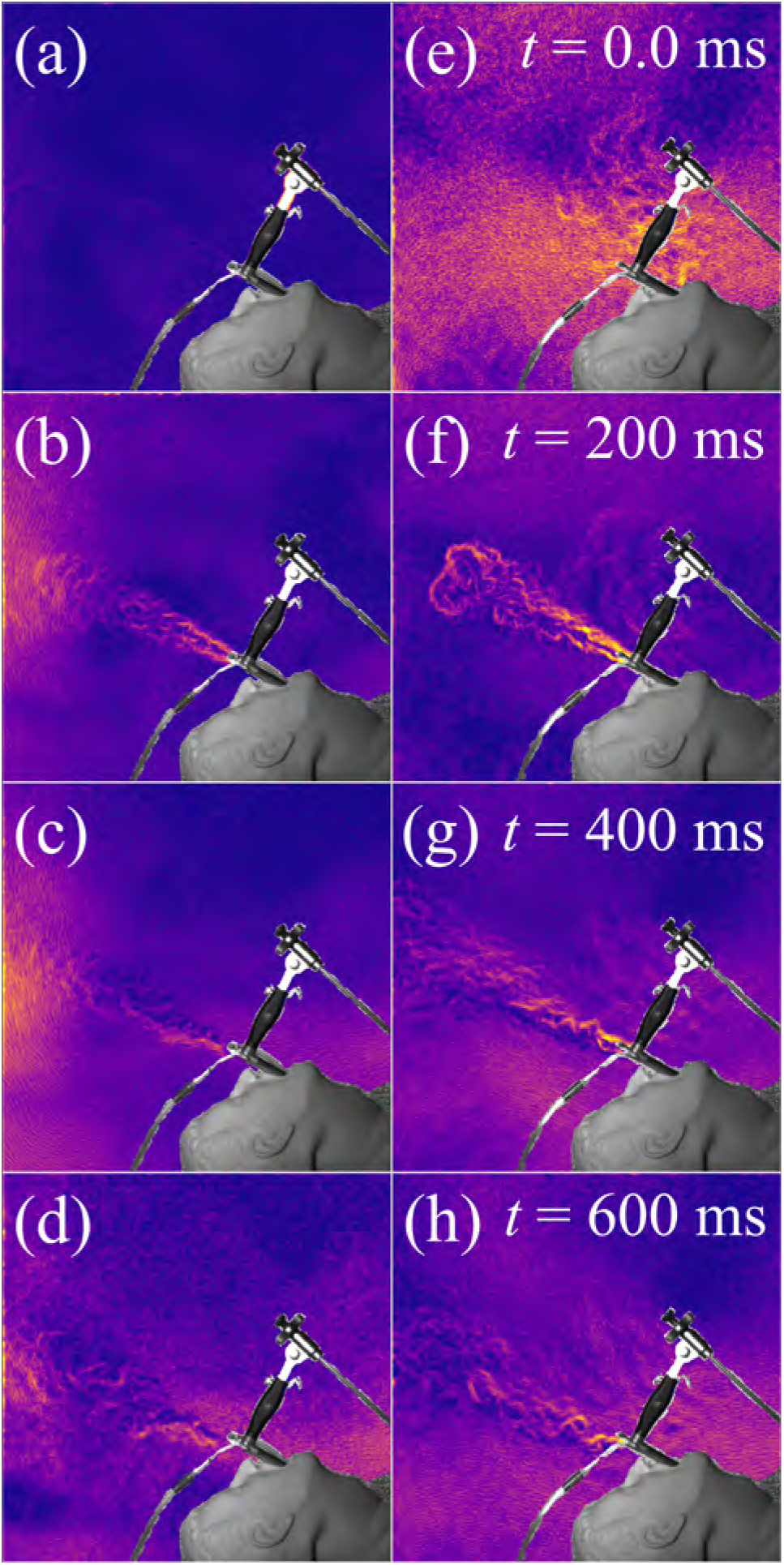
BOS images during jet ventilation. Left column (a)–(d): infraglottic ventilation sequence. Right column (e)–(h) supraglottic ventilation sequence. Note the air issuing from the manikin’s mouth in panel (e).

Images were acquired at 1000 frames per second, and several flow conditions were tested. Properties such as the driving pressure, ventilation frequency, jet ventilation technique, and the use of auxiliary suction. The effects of varying these parameters will be investigated in detail in future work. As an example, Fig. 4 shows BOS-LIC image sequences for two different cases. Both used settings of 30 psi driving pressure, 12 breaths per minute, and suction, but one used infraglottic ventilation (left), in which the ventilation tube is inserted into the trachea, past the larynx, while the other used supraglottic (right), in which the ventilation tube only extends into the laryngoscope, above the larynx. Infraglottic ventilation produces a powerful plume out the laryngoscope as the manikin’s lung compliance pushes air out after the jet ventilator forces air into the lungs. Supraglottic ventilation, on the other hand, produces a weaker flow of air out of the manikin’s mouth *while* the ventilator is forcing air into the lungs with less air projecting from the laryngoscope with a delayed plume from the laryngoscope during the exhalation phase. In both cases the head of the plume travels at approximately 2.4 m/s, which is about 80% faster than plumes created by normal breathing [12]. The plumes are observed to quickly traverse the field of view of 60 × 60 cm, but the maximum distance of propagation could not be determined definitively in our initial cases because of the size of the field of view and camera positioning. It is estimated that the concentrated plumes travel on the order of 30 cm before dispersing to the extent that they are no longer visible with BOS in the present setup.

While continuous auxiliary suction had potential to reduce the amount of aerosol created during the experiments, it was not sufficient to completely eliminate the plumes formed by this AGP. The BOS images suggest a different strategy, wherein the suction is pulsed and synchronized with the jet ventilator, such that suction occurs out of phase with the ventilator. In this way, more powerful suction can be applied that is active only when the aerosol-laden plume would be produced. Experimentation with such a device will be the subject of future work.

### 3.2 Extubation

The same BOS setup was used to evaluate the exhalation from a manikin during simulated extubation. Both intubation and extubation are considered potential AGPs with most current research indicating that extubation is a more significant risk. Extubation involves the removal of the endotracheal tube as a patient emerges from anesthesia. As the patient is waking up, the stimulus of removing the tube is often accompanied by an involuntary cough, which can produce a significant amount of aersol [19]. Within our experiments, the cough was simulated by manually squeezing the manual resuscitator as the intubation tube was removed. A sequence of processed BOS images from this experiment is shown in Fig. 5. The images are not evenly spaced in time, in order to highlight the most important events during the procedure. The plume of exhaled gas is visible issuing from the manikin’s mouth during the simulated cough in the final two frames. The aerosol dispersion can be examined in detail using the video sequence from the high speed camera. The extent of the plume during the extubation is significantly smaller than for jet ventilation, only about 15-20 cm. This is likely due to the lower driving pressure for the cough event. Another factor leading to the decreased plume size is the geometry of the airway from which the plume is produced. In the jet ventilation case, the laryngoscope produces a straight tube about 10 cm in length with a diameter on the order of 1.5 cm, which serves to concentrate the plume and focus its momentum projecting away from the manikin. In the extubation experiment, on the other hand, the plume issues from the manikin’s open mouth and is not formed into a well-defined stream, distributing its momentum over a larger area.

**Fig. 5.**
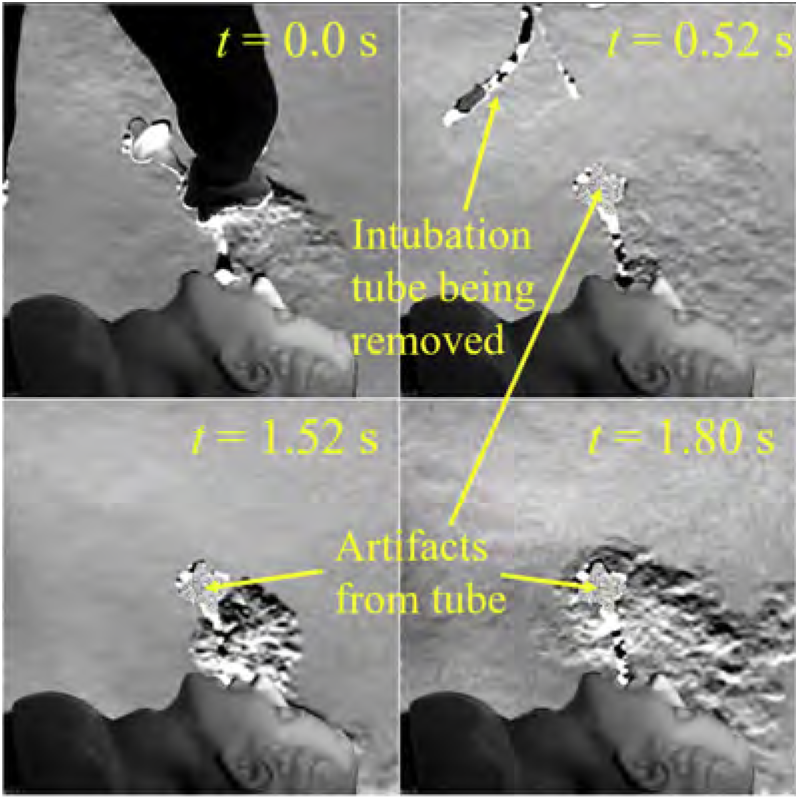
Sequence of BOS images during extubation. Note that the images are not evenly spaced in time, so that the important events could be highlighted. The intubation tube was present in the manikin’s mouth as the reference image was acquired, which is why artifacts from it appear in the final three images after it has been removed.

Unlike in the jet ventilation case where the manikin and laryngoscope are stationary throughout the experiment, in the case of extubation the tube is removed during the experiment. Artifacts are produced by the silhouette of the intubation tube in its initial position in the manikin’s mouth, as shown in Fig. 5. Because the tube is present in the manikin’s mouth when the reference images are acquired, the appearance of the undisturbed background screen behind it cannot be determined, and therefore accurate displacements cannot be calculated there. These artifacts in the displacement field also prevent the use of LIC for visualization, as full streamlines cannot be properly integrated. For AGPs which involve insertion, removal, or repositioning of equipment, therefore, it is recommended that the reference images be acquired with as few objects obscuring the background screen as possible such that the BOS image quality is maximized.

## 4 Discussion

During the COVID-19 pandemic, health care workers deemed to be on the front line were at an increased occupational risk of acquiring the disease despite existing safety protocols. Unprepared medical systems were severely impacted most notably in China, Italy, and several hard-hit U.S. cities. For example, during the time period from February 12—April 9, 2020, 49,370 (16%) of the 315,531 reported COVID-19 cases in the United States were health care workers [20]. Sadly, we had not learned our lesson from previous world pandemics, as this trend was previously seen within severe acute respiratory syndrome (SARS), Ebola, and Middle East respiratory syndrome (MERS) outbreaks [21]. Despite clinical guidelines and hospital best practices formulated by infection control expert recommendations, the limited scientific data, a misunderstanding of the magnitude of the risk and an inability to test potential mitigating policies will ensure future pandemics will have similar deadly impact unless further research can improve our ability to predict and mitigate risk.

It is known that transmission of viral infection can occur through indirect close contact with infected people through both infected respiratory secretions and respiratory droplets [22]. Respiratory droplets, which may include the aerosolized virus, can reach the mucosal surfaces of a susceptible person’s eyes, nose or mouth, resulting in infection. In general, transmission from a patient to a healthcare worker requires that the susceptible person inhale a sufficient quantity of viable virus to cause an infection within the recipient. The exact mechanism of transmission may vary within specific strains of virus or the individual. This aerosolized dose has been studied for other respiratory viruses by measuring the viral RNA via RT-PCR in the cough of symptomatic individuals with Influenza A and B, parainfluenza 1, 2 and 3, respiratory syncytial virus (RSV), and human common cold rhinoviruses (hRV) [23]. Within the Gralton et al. study, the authors confirmed that individuals with symptomatic respiratory viral infections produce both large and small particles carrying viral RNA during coughing and breathing. However, experimental models suggest a high variability between individuals in terms of aerosol emission rates, with increased rates correlated with increased volume of vocalization, for example [24].

### 4.1 Aerosol Generating Procedures

Airborne transmission of infectious particles during medical procedures known as “aerosol generating procedures” (AGPs) are of particular interest. Flow physics studies have generated potential mechanisms of transmission through aerosols [25–28]. Aerosols following specific medical procedures are particularly concerning within healthcare settings in that the aerosols can remain infectious when suspended in the air over long distances and time and may infect susceptible or medically fragile individuals. Indirect contact involving a susceptible host with a contaminated object or surface (known as fomite transmission) may also be possible after respiratory droplets or aerosols contact a surface.

Mitigation techniques may be effective in reducing this risk. Non-physiologic experimental studies using high-powered jet nebulizers under controlled laboratory conditions have demonstrated a relatively low quantity of aerosol detection in an “ultra clean” operating room where air is evacuated into a canopy over the patient’s surgical table [19]. These ultra clean systems perform 500–650 room air exchanges per hour (ACH) through a HEPA filter. By comparison, state building codes require only 15 or 20 ACH in a standard OR. In practice, most hospitals operate at 20 to 25 ACH with some using 30-40 ACH [29]. By comparison, the requirement for patient rooms is 6 ACH. This air exchange rate is important as procedures are performed in a variety of settings and the risk following a AGP is believed to persist for a period of time after procedures, inversely proportionate to the ACH. Typical additional mitigation strategies include recommendations for rooms with 6 ACH to have 60 minutes of down time (non-use time waiting for aerosols to clear) while rooms with 12 ACH require 30 minutes after an AGP.

However, the impact of these types of mitigation strategies are poorly understood. Within the Brown study using a particle counter, a volitional cough as well as intubation and extubation were evaluated, showing that a volitional cough created significantly more aerosol than the act of intubation or extubation [19]. Although post-extubation cough events were noted clinically in about 50% of extubations and the researchers frequently detected an aerosol spike, these spikes were significantly less aerosol-generating than a single volitional cough. Although these extubation related coughs produced a similar particle size distribution, researchers saw 75% fewer airborne particles than during volitional coughs. Thus, the amount of particles generated and the distribution pattern may add considerably more detail to determine the recommended down time after an AGP or potentially reclassify a procedure as a non-AGP. Interestingly, both intubation and extubation may fall below the threshold of a high-risk aerosol-generating procedure with the air exchange rate within the OR having a greater influence on risk [30].

Numerous techniques have been studied in an effort to understand and assist in appropriate classification of medical procedures as AGPs or non-AGPs or would influence down-time between procedures. Methods used include use of particle sensors/optical particle sizers, laser-based particle image velocimetry, continuous air sampling with spectrometry, moisture detection paper, dye studies (e.g. fluorescein isothiocyanate), and surface sampling with reverse transcription quantitative polymerase chain reaction (RT-PCR). In our opinion, this new BOS technique offers a significant advantage of allowing visualization of airflows with actual human patients within a clinical setting that can be used to validate or augment the other listed techniques. Most of these other techniques cannot be performed with actual patients or require that the investigator setup a measurement area which does not necessarily allow the researcher to test the aerosol generation location which may pose the highest risk. For example, in our jet ventilation study, there was a high percentage of airflow emanating from the mouth during supraglottic jet ventilation that would not have been predicted prior to the use of BOS and may enable our development of a number of mitigation strategies.

In the absence of of adequate scientific data to inform us of the true risks, precautionary protective equipment and preventive measures were universally mandated for a majority of hospital systems in response to the COVID-19 pandemic. This has a huge impact on the cost of delivering healthcare and does not focus attention upon the clinical events that place providers at the highest risk. For example, there is a lack of quantitative evidence of the distribution of airborne particles produced during common procedures such as endotracheal intubation to inform risk assessments. Appropriate information would allow infectious disease experts the ability to advise anesthesia providers on appropriate personal protective equipment (PPE) for each type of intervention.

Additional costly protective measures and policies at hospitals include pre-admission laboratory testing on all patients admitted to the hospital as well as elevated PPE protocols for *all* activities involving nearly *all* patients. Early on, many specialty societies recommended rationing of care during the pandemic to reduce healthcare worker exposure to potentially infected patients [31]. This hyper-elevated use of PPE and risk-avoidance posture may unnecessarily utilize resources and exposes patients to complications due to delay of care. Some of the recent studies have called into question these elevated protective measures when pre-operative testing had been performed and whether or not the specific procedure was capable of generating aerosols capable of transmitting an infectious dose [30].

For future viral pandemics, these elevated PPE levels may be unsustainable in outpatient clinics and surgical centers where infection status of the patient is more ambiguous. However, in the absence of knowledge, when an unknown disease state exists, the use of more extensive PPE protocols and avoidance of appropriate care will be performed without good reason. During the current COVID Pandemic this has led to shortages resulting in providers being forced to wear protective gear for extended periods of time reusing soiled masks due to absence of sufficient stock of PPE [32]. It is natural that providers themselves avoid delivery of care deemed high risk when they cannot develop a rational risk assessment. We need

## 5 Conclusion

AGPs include a variety of routine health care procedures and patient care activities that have a variable potential of transmission of airborne pathogens. The technique described here adapts BOS visualization of air flows from relatively small-scale experimental fluid mechanics problems, to a highly portable and low cost system capable of visualizing fields of view on the order of 100 cm and larger in each dimension. The work presented here represents a demonstration of the capabilities of the system only, but importantly, it lays the groundwork for in-depth analysis of AGPs in both simulated and clinical studies using BOS and the development of risk mitigation strategies, such as the synchronous suction suggested in Sec. 3.1. In addition, this work features the first application of LIC to BOS images to the best of the authors’ knowledge, which allows for compelling, intuitive visualization of both components of the two-dimensional displacement field along with its magnitude. Future clinical use of this technique will require achieving substantially larger fields of view of 2-3 m and allow for intermittent disruptions in the camera view of the background. These larger fields of view will aid in the determination of risk and the subsequent management of aerosol generating procedures within a clinical space. Understanding and managing the risk is a critical step to help our healthcare team respond to this and future pandemics.

## Acknowledgements

N. S. H. and B. E. S. acknowledge National Institute of Biomedical Imaging and Bioengineering (NIBIB) 1-R21-EB032644-01, which supported this study.

## References

[1] Klompas, M., Baker, M., and Rhee, C., 2020. “What is an aerosol-generating procedure?”. JAMA Surgery.

[2] Jackson, T., Deibert, D., Wyatt, G., Durand-Moreau, Q., Adisesh, A., Khunti, K., Khunti, S., Smith, S., Chan, X. H. S., Ross, L., Roberts, N., Toomey, E., Greenhalgh, T., Arora, I., Black, S. M., Drake, J., Syam, N., Temple, R., and Straube, S., 2020. “Classification of aerosol-generating procedures: a rapid systematic review”. BMJ Open Respiratory Research, 7(1).

[3] Bourouiba, L., 2020. “Turbulent gas clouds and respiratory pathogen emissions”. JAMA.

[4] Bahl, P., Doolan, C., de Silva, C., Chughtai, A. A., Bourouiba, L., and MacIntyre, C. R., 2022. “Airborne or droplet precautions for health workers treating coronavirus disease 2019?”. The Journal of Infectious Diseases, 225(9), pp. 1561–1568.

[5] Raffel, M., Willert, C. E., Scarano, F., Kähler, C. J., Wereley, S. T., and Kompenhans, J., 2018. Particle image velocimetry: a practical guide. Springer.

[6] Poggie, J., Erbland, P. J., Smits, A. J., and Miles, R. B., 2004. “Quantitative visualization of compressible turbulent shear flows using condensate-enhanced Rayleigh scattering”. Experiments in Fluids, 37(3), pp. 438–454.

[7] Settles, G. S., 2001. Schlieren and Shadowgraph Techniques, first ed. Springer Berlin Heidelberg.

[8] Tang, J. W., Liebner, T. J., Craven, B. A., and Settles, G. S., 2009. “A schlieren optical study of the human cough with and without wearing masks for aerosol infection control”. Journal of The Royal Society Interface, 6(suppl_6).

[9] Tang, J. W., Nicolle, A. D. G., Pantelic, J., Jiang, M., Sekhr, C., Cheong, D. K. W., and Tham, K. W., 2011. “Qualitative real-time schlieren and shadowgraph imaging of human exhaled airflows: An aid to aerosol infection control”. PLoS ONE, 6(6).

[10] Staymates, M., 2020. “Flow visualization of an N95 respirator with and without an exhalation valve using schlieren imaging and light scattering”. Physics of Fluids, 32(11).

[11] Raffel, M., 2015. “Background-oriented schlieren (BOS) techniques”. Experiments in Fluids, 56(3).

[12] Mok, T., Harris, E., Vargas, A., Afshar, Y., Han, C. S., Karagozian, A., and Rao, R., 2021. “Evaluation of respiratory emissions during labor and delivery”. Obstetrics & Gynecology, 138(4), pp. 616–621.

[13] Schmidt, B. E., and Woike, M. R., 2021. “Wavelet-based optical flow analysis (wOFA) for background oriented schlieren (BOS) image processing”. AIAA Journal.

[14] Laidlaw, D. H., Kirby, R. M., Jackson, C. D., Davidson, J. S., Miller, T. S., da Silva, M., Warren, W. H., and Tarr, M. J., 2005. “Comparing 2D vector field visualization methods: A user study”. IEEE Transactions on Visualization and Computer Graphics, 11(01), pp. 59–70.

[15] Cabral, B., and Leedom, L. C., 1993. “Imaging vector fields using line integral convolution”. In Proceedings of the 20th annual conference on computer graphics and interactive techniques, pp. 263–270.

[16] Atcheson, B., Heidrich, W., and Ihrke, I., 2009. “An evaluation of optical flow algorithms for background oriented schlieren imaging”. Experiments in Fluids, 46(3), pp. 467–476.

[17] Cook, R. L., and DeRose, T., 2005. “Wavelet noise”. ACM Transactions on Graphics, 24(3), pp. 803–811.

[18] Crameri, F., Shephard, G. E., and Heron, P. J., 2020. “The misuse of colour in science communication”. Nature Communications, 11(1).

[19] Brown, J., Gregson, F. K. A., Shrimpton, A., Cook, T. M., Bzdek, B. R., Reid, J. P., and Pickering, A. E., 2020. “A quantitative evaluation of aerosol generation during tracheal intubation and extubation”. Anaesthesia, 76(2), pp. 174–181.

[20] Burrer, S. L., de Perio, M. A., Hughes, M. M., Kuhar, D. T., Luckhaupt, S. E., McDaniel, C. J., Porter, R. M., Silk, B., Stuckey, M. J., and Walters, M., 2020. “Characteristics of health care personnel with COVID-19 — united states, february 12–april 9, 2020”. MMWR. Morbidity and Mortality Weekly Report, 69(15), pp. 477–481.

[21] Knoetze, C., 2020. Protecting the health workers who protect us all. Online, Sept.

[22] Liu, J., Liao, X., Qian, S., Yuan, J., Wang, F., Liu, Y., Wang, Z., Wang, F., Liu, L., and Zhang, Z., 2020. “Community transmission of severe acute respiratory syndrome coronavirus 2, Shenzhen, China, 2020”. Emerging Infectious Diseases, 26(6).

[23] Gralton, J., Tovey, E. R., McLaws, M.-L., and Rawlinson, W. D., 2013. “Respiratory virus RNA is detectable in airborne and droplet particles”. Journal of Medical Virology, 85(12), pp. 2151–2159.

[24] Asadi, S., Wexler, A. S., Cappa, C. D., Barreda, S., Bouvier, N. M., and Ristenpart, W. D., 2019. “Aerosol emission and superemission during human speech increase with voice loudness”. Scientific Reports, 9(1).

[25] Fowler, R. A., Guest, C. B., Lapinsky, S. E., Sibbald, W. J., Louie, M., Tang, P., Simor, A. E., and Stewart, T. E., 2004. “Transmission of severe acute respiratory syndrome during intubation and mechanical ventilation”. American Journal of Respiratory and Critical Care Medicine, 169(11), pp. 1198–1202.

[26] Simonds, A. K., Hanak, A., Chatwin, M., Morrell, M. J., Hall, A., Parker, K. H., Siggers, J. H., and Dickinson, R. J., 2010. “Evaluation of droplet dispersion during non-invasive ventilation, oxygen therapy, nebuliser treatment and chest physiotherapy in clinical practice: implications for management of pandemic influenza and other airborne infections”. Health Technology Assessment, 14(46).

[27] Esquinas, A. M., Pravinkumar, S. E., Scala, R., Gay, P., Soroksky, A., Girault, C., Han, F., Hui, D. S., Papadakos, P. J., and Ambrosino, N., 2014. “Noninvasive mechanical ventilation in high-risk pulmonary infections: a clinical review”. European Respiratory Review, 23(134), pp. 427–438.

[28] Thille, A. W., Muller, G., Gacouin, A., Coudroy, R., Decavèle, M., Sonneville, R., Beloncle, F., Girault, C., Dangers, L., Lautrette, A., Cabasson, S., Rouzé, A., Vivier, E., Le Meur, A., Ricard, J., Razazi, K., Barberet, G., Lebert, C., Ehrmann, S., Sabatier, C., Bourenne, J., Pradel, G., Bailly, P., Terzi, N., Dellamonica, J., Lacave, G., Danin, P., Nanadoumgar, H., Gibelin, A., Zanre, L., Deye, N., Demoule, A., Maamar, A., Nay, M., Robert, R., Ragot, S., and Frat, J., 2019. “Effect of postextubation high-flow nasal oxygen with noninvasive ventilation vs high-flow nasal oxygen alone on reintubation among patients at high risk of extubation failure”. JAMA, 322(15), p. 1465.

[29] Gormley, T., and Wagner, J., 2018. Studying airflow in the OR: measuring the environmental quality indicators in a dynamic hospital operating room setting. Online, Jan.

[30] White, S. M., and Chakladar, A., 2020. “Aerosols: are anaesthetists at risk of COVID-19 or not?”. Anaesthesia.

[31] Gralnek, I. M., Hassan, C., Beilenhoff, U., Antonelli, G., Ebigbo, A., Pellisè, M., Arvanitakis, M., Bhandari, P., Bisschops, R., Van Hooft, J. E., Kaminski, M. F., Triantafyllou, K., Webster, G., Pohl, H., Dunkley, I., Fehrke, B., Gazic, M., Gjergek, T., Maasen, S., Waagenes, W., de Pater, M., Ponchon, T., Siersema, P. D., Messmann, H., and Dinis-Ribeiro, M., 2020. “ESGE and ESGENA position statement on gastrointestinal endoscopy and the COVID-19 pandemic”. Endoscopy, 52(06), pp. 483–490.

[32] Gralnek, I. M., Hassan, C., Beilenhoff, U., Antonelli, G., Ebigbo, A., Pellise, M., Arvanitakis, M., Bhandari, P., Bisschops, R., Van Hooft, J. E., Kaminski, M. F., Triantafyllou, K., Webster, G., Pohl, H., Dunkley, I., Fehrke, B., Gazic, M., Gjergek, T., Maasen, S., Waagenes, W., de Pater, M., Ponchon, T., Siersema, P. D., Messmann, H., and Dinis-Ribeiro, M., 2020. Safety of extended use and reuse of N95 respirators. ECRI Clinical Evidence Assessment, Mar.

